# Impact of aldehyde dehydrogenase 2 deficiency on tissue-specific mitochondrial metabolism in aging mice

**DOI:** 10.64898/2026.05.25.727652

**Authors:** Marcio Augusto Campos-Ribeiro, Vanessa Morais Lima, Thiago N. Menezes, Wenjin Yang, Juliane Cruz Campos, Julio Cesar Batista Ferreira

## Abstract

Age-related diseases arise from prolonged exposure to genetic and/or environmental factors, ultimately leading to cumulative and irreversible degeneration of tissues and the organism as a whole. We previously reported that accumulation of mitochondrially-generated aldehydes (i.e., 4-hydroxynonenal and acetaldehyde) causes mitochondrial dysfunction and accelerates the progression of age-related diseases. However, the contribution of mitochondrial aldehyde metabolism to aging (via aldehyde dehydrogenase 2, ALDH2) remains elusive. Here, we provide a comprehensive analysis of aldehyde metabolism and mitochondrial bioenergetics across different tissues in aging mice. We also address how mitochondrial function is influenced by the highly prevalent human inactivating ALDH2 E504K point mutation (ALDH2^E504K^) during aging. The liver metabolism was relatively resilient to aging, showing enhanced ALDH2 activity and improved mitochondrial coupling. Strikingly, aging-associated liver resilience was lost in ALDH2^E504K^ mice. Aged hearts exhibited mixed outcomes including impaired mitochondrial basal respiration, improved ADP-driven respiration, and decreased ALDH2 detox capacity. The ALDH2^E504K^ mutation exacerbated the already impaired cardiac ALDH2 detox capacity in aging. Strikingly, aging brain displayed pronounced vulnerability, with decreased ALDH2 activity, impaired mitochondrial bioenergetics and defective ALDH2 detox capacity. These changes were paralleled by impaired cognitive and behavioral functions in aged mice. As proof of concept, either the presence of ALDH2^E504K^ mutation or acute ethanol challenge worsened cognitive and behavioral dysfunction in aging mice. Finally, we assessed *in vitro* efficacy of pharmacological ALDH2 activation in aging tissues. Collectively, these findings unravel the contribution of ALDH2^E504K^ mutation to mitochondrial metabolism during aging; highlighting the detrimental synergy between genetic ALDH2 deficiency and aging in brain metabolism and physiology.

## 1. INTRODUCTION

Age-related disorders are (un)coordinated, multifaceted and degenerative processes affected by both genetic and environmental factors. They have become more prevalent worldwide as result of unhealthy living habits and increased life expectancy (1). Therefore, the precise understanding of critical biological effectors of age-related diseases is required to develop better interventions capable of promoting healthy aging.

Impairment of mitochondrial detoxification functions is pivotal in the establishment and progression of age-related disorders (2, 3). Among different damaging reactive molecules accumulated in aging (4), mitochondrially-derived aldehydes produced during several intracellular processes including (but not limited to) mitochondrial oxidative stress (i.e., 4-hydroxynonenal and malondialdehyde (5), mitochondrial glucose metabolism (i.e., acetaldehyde (6)), and mitochondrial ethanol metabolism (i.e., acetaldehyde (7)) are extremely harmful. These aldehydes contribute to age-related diseases by forming adducts with macromolecules including proteins, DNA and lipids; therefore, inhibiting and/or disrupting their function (8-11). As consequence, aldehydic load impairs essential biological processes such as DNA replication (12), DNA transcription (12), mitochondrial metabolism (5) and miRNA biogenesis (13).

We have previously reported that pharmacological activation of aldehyde dehydrogenase 2 (ALDH2), a mitochondrial enzyme responsible for oxidizing reactive aldehydes, is sufficient to protect against age-related diseases including myocardial infarction (14) and heart failure (5). ALDH2 protection occurs, at least in part, through improving local clearance of 4-hydroxynonenal (14) and acetaldehyde (7). As expected, loss of ALDH2 function maximizes cardiac damage caused by accumulation of mitochondrially-derived reactive aldehydes (7); therefore, strengthen the role of ALDH2 as a potential target to treat age-related disorders.

Despite its critical role in several animal models of age-related diseases, the contribution of ALDH2 to aging remains elusive. Here, we used genetically engineered knock-in animals to study the role of ALDH2 in aging. These mice carry a single amino acid substitution that is equivalent to the E504K substitution that defines the human ALDH2^E504K^ mutation (15). This inactivating point mutation results in a dramatic decline in ALDH2 activity in both rodents (15) and humans (16). Note that about 560 million East Asians (8% of the world population) carry this ALDH2^E504K^ mutation (17), which is probably the most common human enzyme deficiency worldwide. The ALDH2^E504K^ mutation has been recently associated with a variety of age-related disorders including cancer (18), cardiovascular diseases (19), Alzheimer disease (20), and skeletal muscle weakness (21) in humans. However, its contribution to tissue-specific function, metabolism and bioenergetics in aging remains to be determined.

Here, we set out to determine the contribution of ALDH2 deficiency to tissue-specific aldehyde detoxification and mitochondrial function in aging mice. For that, we assessed both levels and activity of ALDH2, and mitochondrial bioenergetics in different tissues from young (3-month-old) and old (24-month-old) wild-type (WT), ALDH2^E504K^ heterozygous and homozygous mice. Considering the central role of ALDH2 on ethanol metabolism, we also performed some behavioral tests in ALDH2 deficient young and aged animals acutely challenged with ethanol. Finally, we tested the responsiveness of different tissues from young and old animals to selective pharmacological activator (AD-5591) and inhibitor (cvt-10216) of ALDH2.

## 2 MATERIALS AND METHODS

### 2.1 Ethics approval

This study was conducted in accordance with the ethical principles in animal research adopted by the Brazilian College of Animal Experimentation (www.cobea.org.br). The animal care and protocols in this study were reviewed and approved by the Ethical Committee of the Institute of Biomedical Sciences at University of São Paulo (#133123).

### 2.2 Animal care and use

A cohort of 3- and 24-month-old male wild-type, ALDH2^E504K^ heterozygous and homozygous C57BL/6J mice was selected for the study. Knock-in ALDH2^E504K^ mouse carrying the human inactivating ALDH2 point mutation E504K with a C57BL/6J background were generated by homologous recombination as described elsewhere (15). Knock-in ALDH2^E504K^ mice are an ideal representation of the human ALDH2^E504K^ carriers and can serve as an experimental model to reflect the true nature of the genetic defect, the ALDH2 deficiency. Animals were maintained in a 12:12-h light-dark cycle and temperature-controlled environment (22°C) with free access to standard laboratory chow (Nuvital Nutrientes, Curitiba, PR, Brazil) and tap water. Body weight was measured weekly. For the acute ethanol challenge, animals received either water or ethanol (2g/kg) by gavage, and 180 minutes later behavior experiments were performed. At the end of experimental protocol, animals were weighted, anesthetized (isoflurane, 5%, flow rate 0.5 l/min) and killed by cervical dislocation to harvest their tissues for biochemical measurements. Excised tissues were weighted, processed and stored at −80C.

### 2.3 Open field test

To evaluate locomotor activity and vertical exploratory ability; animals were submitted to the open field test. The open field test consisted of a circular arena with three concentric circles. The outer ring was divided into 16 parts, the inner one into 8 parts, and the central one had no divisions. During the test, each animal was placed in the center of the arena and allowed to explore the environment for 5 minutes. During this period, the number of quadrants crossed by the animal (locomotor activity) and the number of times the animal reared on its hind legs to explore the environment were analyzed. The mouse activity was digitally recorded using a video camera placed above the center of the arena. At the end of each test the arena was cleaned with 70% (v/v) ethanol to prevent odor cues.

### 2.4 Novel object recognition test

To assess memory and learning; animals were subjected to the novel object recognition test. The test consisted of a sample and a test phase. During sample phase, animals from each group were individually placed into a box containing two identical objects. Animals were allowed to explore the box and the objects for 5 minutes. After 3 hours, one familiar object in the box was replaced with a different object (novel object) and the test phase began. Each animal was recorded for 5 minutes while exploring the objects (novel and familiar). Exploration activity was considered when the animal pointed its snout at the object. In both sample and test phases the amount of time each animal spent exploring the two objects was considered. In the training stage, it is expected that the animals explore both objects equally. In the test stage, the animal is expected to explore more the novel object, to the detriment of the familiar object. The discrimination index was calculated using the following formula: discrimination index = (time with novel object – time with familiar object)/ (time with novel object + time with familiar object). After each run, objects and boxes were cleaned with 70% (v/v) ethanol to prevent odor cues.

### 2.5 Elevated plus maze test

The elevated plus maze, which is used to assess animals’ anxiety, consists of 2 opposite open arms (50 cm x 10 cm) and 2 closed arms, also opposite, in the shape of a plus sign, 50 cm above the floor. The animals were placed at the intersection between the open and closed arms, and left to explore the apparatus for 5 minutes. Anxiety was calculated by measuring the percentage of time animals spent in the open arms compared to the closed arms. At the end of each test the maze was sanitized with 70% alcohol.

### 2.6 Y-maze test

The Y maze test was performed to assess spatial working memory and exploratory behavior. Briefly, animals were placed at the end of one arm of a Y-shaped maze with 3 identical arms (35 cm long, 8 cm wide, named A, B, and C), and then allowed to freely explore all three arms for 5 minutes. Arm entries were recorded using an automated video tracking system and confirmed by manual scoring. An arm entry was defined as the placement of all four paws into the arm. The primary measure was spontaneous alternation behavior, reflecting the natural tendency of rodents to enter a less recently visited arm. Alternation was defined as successive entries into all three different arms (e.g., ABC, BCA, CAB). The percentage of spontaneous alternation was calculated using the formula: % alternation=(number of alternations/total arm entries-2)X100. At the end of each test the maze was sanitized with 70% alcohol.

### 2.7 Hindlimb clasping test

The hindlimb clasping test was performed to assess motor impairment or neurological dysfunction. Briefly, each mouse was suspended by the tail for 10 seconds in a quiet room under constant illumination. During the tail-suspension, a score was assigned to each animal according to the positioning of its paws. The scores assigned were: 0 - animal positioned its hind legs open away from the abdomen and fingers open; 1 - animal had one of its legs with incomplete opening and fingers opened; 2 - animal had both hind legs with incomplete opening and fingers opened; 3 - animal had its hind legs close to the abdomen and attached to each other with closed fingers; and 4 - animal had its front and hind legs crossed, fingers closed and immobile (22).

### 2.8 Rotarod test

The rotarod test was used to assess motor coordination and balance. Briefly, animals were placed on a rotarod apparatus consisting of a rotating rod (3 cm in diameter, 30 cm in length) divided into multiple lanes to test several mice simultaneously. The rod was elevated 25 cm above a padded surface to prevent injury in case of falls. Mice were subjected to a training session one day before testing to familiarize them with the apparatus. During the test, each animal was placed individually on the rotating rod, which accelerated from 4 to 40 revolutions per minute (rpm) over 3 minutes. The trial ended when the mouse fell from the rod or after a maximum cutoff time of 300 seconds. Each animal underwent three consecutive trials separated by 15-minute inter-trial intervals to avoid fatigue. The latency to fall (in seconds) was recorded automatically by the apparatus. The average latency to fall across trials was calculated for each animal and used as the primary index of motor coordination and balance. A longer latency to fall indicates improved motor performance, while a shorter latency reflects motor impairment.

### 2.9 Voluntary wheel running test

Voluntary wheel running activity was measured by individually housing mice in cages equipped with low-resistance running wheels connected to digital counters. Wheel revolutions were continuously recorded for 14 days and converted into running distance (km) using the wheel circumference. Running activity was expressed as running distance, average speed and running time. Food and water were provided ad libitum.

### 2.10 Mitochondrial isolation

Cardiac, liver and brain mitochondria were isolated as described elsewhere (5). Briefly, samples were minced and homogenized in isolation buffer (300 mM sucrose, 10 mM HEPES, 2 mM EGTA, pH 7.2, 4°C). The suspension was homogenized in a 4 mL tissue grinder and centrifuged at 950 g for 5 min. The resulting supernatant was centrifuged at 9500 g for 10 min. The mitochondrial pellet was washed, resuspended in isolation buffer, and submitted to a new centrifugation (9500 g for 10 min). The mitochondrial pellet was washed, and the final pellet was resuspended in a minimal volume of isolation buffer.

### 2.11 Assessment of mitochondrial H_2_O_2_ release and O_2_ consumption

Mitochondrial H_2_O_2_ release and O_2_ consumption measurements were performed as described elsewhere (5). Briefly, mitochondrial H_2_O_2_ release was determined by measuring the oxidation of Amplex Red in the presence of horseradish peroxidase using a spectrophotometer with 563 nm of excitation and 587 nm of emission. Mitochondrial O_2_ consumption was monitored using a computer-interfaced Clark-type electrode (OROBOROS, Oxygraph-2k). Experiments were carried out in buffer containing 125 mM sucrose, 65 mM KCl, 10 mM Hepes, 2 mM KH_2_PO_4_, 2 mM MgCl_2_, 0.01% BSA, pH 7.2; and 0.125 mg/mL protein. Measurements were acquired under continuous stirring at 37°C. Succinate, malate, and glutamate (2 mmol/L of each) were used as substrates, and ADP (1 mmol/L) was added to induce State 2 and State 3 respiratory rates, respectively.

### 2.12 Immunoblotting

ALDH2 protein levels were evaluated in different tissues from young and old mice as previously described (13). Briefly, samples were subjected to SDS-PAGE and proteins were electro transferred onto nitrocellulose membranes (Bio-Rad Biosciences, Piscataway, NJ, USA). Equal gel loading and transfer efficiency were monitored using 0.5% Ponceau S staining of blot membrane. Blotted membranes were then blocked [5% non-fat dry milk, 10 mM Tris–HCl (pH 7.6), 150 mM NaCl, and 0.1% Tween 20] for 2 h at room temperature and then incubated overnight at 4°C with specific antibodies. Binding of the primary antibody was detected with the use of peroxidase-conjugated secondary antibodies (rabbit or mouse, depending on the protein, for 2 h at room temperature) and developed using enhanced chemiluminescence (Amersham Biosciences, NJ, USA) detected by autoradiography. Quantification analysis of blots was performed with the use of Scion Image software (Scion based on NIH image). Samples were normalized to relative changes in housekeeping (VDAC or ponceau).

### 2.13 ALDH2 enzyme activity

Enzymatic activity of ALDH2 was determined by measuring the conversion of NAD^+^ to NADH, as described elsewhere (23). The assays were carried out at 25°C in 50 mM sodium pyrophosphate buffer (pH 9.5) in the presence of 10 mM acetaldehyde. Measurement of ALDH2 activity was determined by directly adding 80 μg of the mitochondrial fraction of tissue to the reaction mix and reading absorbance at 340 nm for 10 min. For the *in vitro* study measuring the effect of AD-5591 (ALDH2 activator) and cvt-10216 (ALDH2 inhibitor) on ALDH2 activity from different tissues, lysates were incubated with either AD-5591 (1μM) or cvt-10216 (1μM) 5 minutes prior to the addition of ALDH2 substrate acetaldehyde. To estimate the relationship between mitochondrial aldehyde-detoxifying activity and oxidative burden, we calculated the ALDH2 detox capacity index, a ratio between ALDH2 activity (a player involved in aldehydic unload) and state 2 mitochondrial hydrogen peroxide release (a player involved in aldehydic overload).

### 2.14 Statistical Analysis

Data are presented as means ± standard error of the mean (SEM). Data normality was assessed through Shapiro–Wilk test. Group differences were calculated using two-tailed Student’s t-test for pairwise comparisons and two-way analysis of variance (ANOVA) for multiple comparisons involving two independent variables. Whenever significant F-values were obtained, Tukey adjustment was used for multiple comparison purposes. To integrate ALDH2 metabolism and mitochondrial bioenergetics variables across tissues and age groups, we employed a combination of principal component analysis (PCA) and permutational multivariate analysis of variance (PERMANOVA). GraphPad Prism Statistics (version 11; GraphPad Software, Inc., La Jolla, CA) was used for the analysis, and statistical significance was considered achieved when the *P*-value was < 0.05. Raw data are presented as individual data points in the graphs.

## 3. RESULTS

### 3.1 Tissue-specific ALDH2 metabolism and mitochondrial bioenergetics are differently affected by aging

ALDH2 plays a beneficial role in several animal models of age-related degenerative diseases (5, 14, 24, 25). However, the proposed loss of ALDH2 function in aging remains to be determined. Here, we first assessed ALDH2 protein abundance across different tissues from young (3-mo-old) and aged (24-mo-old) wild-type (WT) mice (Fig. 1A). Liver, lung, kidney and heart tissues display elevated ALDH2 protein levels compared with skeletal muscle (plantaris) and brain at both ages (Fig. 1B). During aging, there is no major change in ALDH2 protein levels in liver, lung, kidney, heart and skeletal muscle (plantaris) (Fig. 1B). Surprisingly, ALDH2 protein levels were significantly elevated by 80 ± 30% in brain tissue from aged mice (Fig. 1B). To further explore mitochondrial function in aging, we measured both ALDH2 catalytic activity and mitochondrial bioenergetics in liver, heart and brain. We have chosen these tissues because liver is the primary organ for aldehyde detoxification (26), and heart and brain display elevated baseline energy demand (27). We have previously reported that ALDH2 dysfunction contributes to impaired mitochondrial bioenergetics and progression of aging-related degenerative disorders such as metabolic, cardiovascular and neurological diseases (5, 14, 24, 25).

**Fig. 1.**
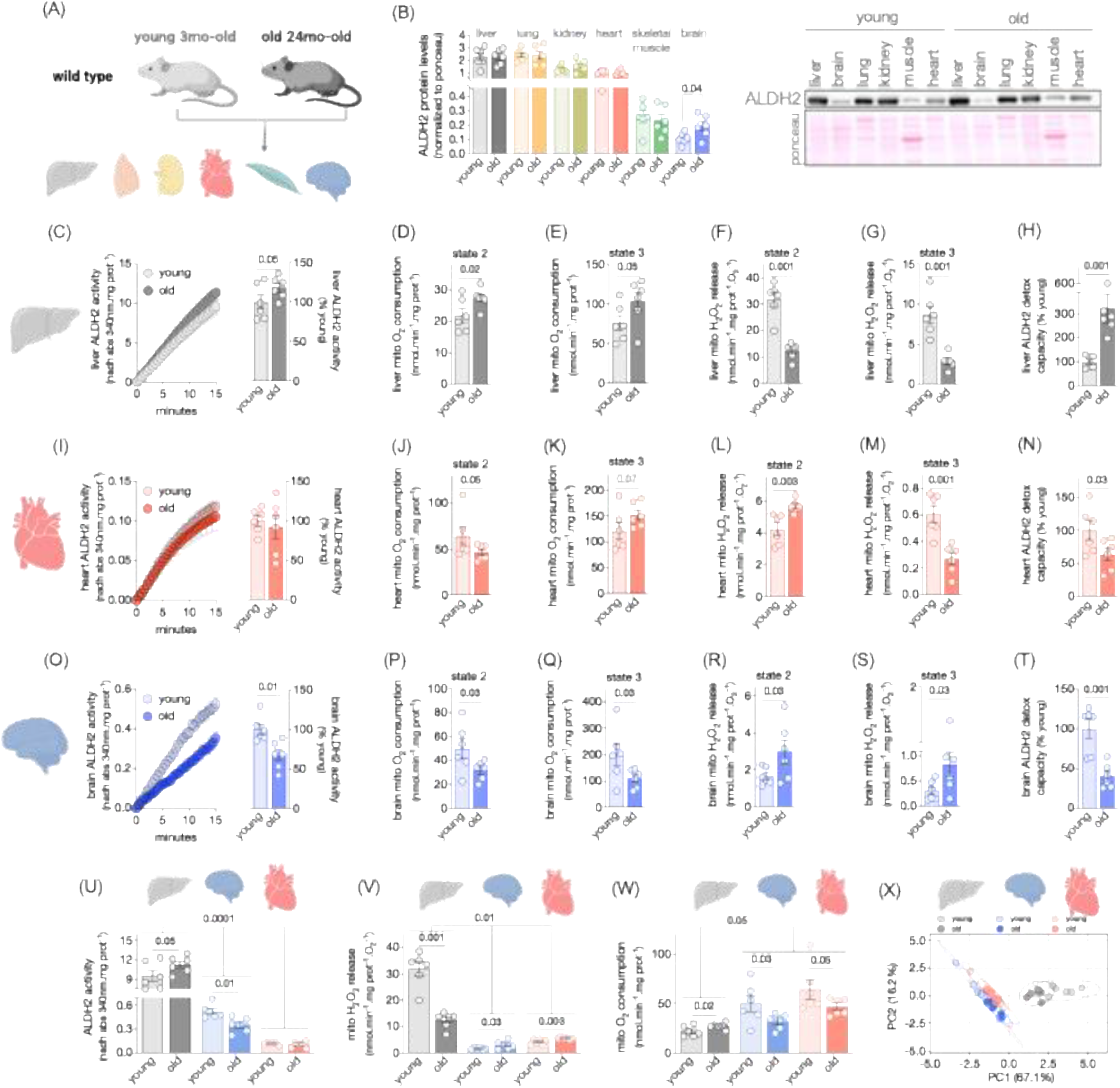
Tissue-specific profile of ALDH2 activity and mitochondrial bioenergetics during aging. (A) Schematic panel - mitochondrial metabolism was measured in different tissues from young (3-month-old, n=6) and old (24-month-old, n=6) male C57BL/6J mice. (B) ALDH2 protein levels in liver, lung, kidney, heart, skeletal muscle, and brain (left panel: quantification - individual data and average of all samples, normalized by ponceau staining; right panel: representative ALDH2 immunoblot and ponceau staining). (C) ALDH2 activity (left panel: average slope; right panel: quantification - individual data and average of all samples, expressed as % of young mice), (D–E) mitochondrial O_2_ consumption and (F–G) mitochondrial H_2_O_2_ release (expressed as H_2_O_2_/O_2_) measured in isolated mitochondria under state 2 (after addition of succinate 10 mM, malate 2 mM, glutamate 10 mM) and state 3 (after addition of ADP 4 mM) respiratory rates, and (H) ALDH2 detox capacity (expressed as ALDH2 activity divided by mitochondrial H_2_O_2_ release) in liver (C-H), heart (I-N) and brain (O-T) tissues. (U) ALDH2 activity, (V) state 2 mitochondrial H_2_O_2_ release, (W) state 2 mitochondrial O_2_ consumption, and (X) Principal component analysis (PCA) of ALDH2 activity and mitochondrial bioenergetics variables in liver, brain, and heart from young (3-month-old) and old (24-month-old) wild-type mice (n=6 per group). To estimate the relationship between mitochondrial aldehyde-detoxifying activity and oxidative burden, we calculated the ALDH2 detox capacity index, a ratio between ALDH2 activity (a player involved in aldehydic unload) and state 2 mitochondrial hydrogen peroxide release (a player involved in aldehydic overload). Data are presented as individual values and means ± SEM. Statistical significance (p<0.05) was assessed using two-tailed Student’s t-tests for pairwise comparisons (B–T), and two-way analysis of variance (ANOVA) followed by Tukey’s post-hoc test (U–W). P values are individually shown in each panel. Permutational multivariate analysis of variance (PERMANOVA) was used to determine significance in the PCA (X).

ALDH2 activity was significantly elevated in liver tissue from aged mice compared with young animals (Fig. 1C). Of interest, such improvement in ALDH2 activity was paralleled by increased mitochondrial oxygen consumption in aged animals (Fig. 1D-E). Old mice displayed higher mitochondrial state 2 (basal) and state 3 (ADP-induced respiration) respiratory rates compared with young animals (Fig. 1D-E). Elevated baseline mitochondrial oxygen consumption has been associated with reactive oxygen species production during mitochondrial uncoupling in aging-related diseases (28). To further test the interdependence of mitochondrial oxygen consumption and reactive oxygen species production, we assessed mitochondrial hydrogen peroxide release. Liver mitochondria isolated from aged animals displayed decreased state 2 and state 3 hydrogen peroxide release compared with young mice (Fig. 1F-G). Note that mitochondrial hydrogen peroxide release was normalized by oxygen consumption rates, demonstrating a significant decrease of hydrogen peroxide release per molecule of oxygen consumed in aging liver. Finally, liver ALDH2 detox capacity (expressed as ALDH2 catalytic activity divided by mitochondrial hydrogen peroxide release) was elevated in aged versus young mice (Fig. 1H). These findings suggest that liver mitochondria isolated from aged mice display improved aldehyde metabolism and bioenergetics compared with young animals.

We next measured ALDH2 activity and mitochondrial bioenergetics in cardiac tissue. Aged mice displayed no changes in cardiac ALDH2 activity but decreased state 2 (basal) mitochondrial respiratory rates and elevated hydrogen peroxide release compared with young animals (Fig. 1I-J and L). These findings suggest impaired mitochondrial bioenergetics in aging hearts. Intriguingly, old mice exhibited a trend toward increased ADP-coupled mitochondrial respiratory rates and decreased hydrogen peroxide release compared with young animals (Fig. 1K and M), indicating improved coupling of oxygen consumption with oxidative phosphorylation. Cardiac ALDH2 detox capacity was significantly decreased in aged versus young animals (Fig. 1N).

Finally, we assessed mitochondrial function in brain tissue. Surprisingly, aging brain displayed a distinct metabolic profile compared with liver and heart tissues. There was a significant decrease in brain ALDH2 activity in aged versus young mice (Fig. 1O). Such impairment in ALDH2 activity was paralleled by a decline in mitochondrial respiratory state 2 and state 3 rates, and exacerbated hydrogen peroxide release in aging brains (Fig. 1P-S). As consequence, the ALDH2 detox capacity was impaired in brains from old versus young mice (Fig. 1T).

In summary, liver, heart and brain tissues displayed heterogeneous mitochondrial ALDH2 metabolism and bioenergetics profile during aging. Liver tissue exhibited the highest ALDH2 activity and mitochondrial peroxide release per molecule of oxygen consumed, despite having the lowest mitochondrial oxygen consumption, compared with heart and brain tissues (Fig. 1U-W and S1A-B). Intriguingly, liver tissue was more resistant while heart and brain were more susceptible to aldehydic load and mitochondrial bioenergetics dysfunction during aging (Fig. 1U-W). Indeed, the principal component analysis (PCA) revealed a strong tissue-driven separation (R^2^ = 0.77, p = 0.001), which was not associated with differences in dispersion, indicating a robust tissue effect (Fig. 1X and S1C). Age-related differences were secondary and tissue-dependent. While liver showed a significant age effect (R^2^ = 0.59, p = 0.002), substantial overlap was observed in heart (R^2^ = 0.21, p = 0.066) and brain (R^2^ = 0.29, p = 0.052). These findings suggest that tissues with higher bioenergetics demand (i.e., heart and brain) are more prone to aging-induced aldehydic load and mitochondrial dysfunction.

### 3.2 Liver ALDH2 buffering capacity is impaired in aged ALDH2 deficient mice

Considering that ALDH2 activity and mitochondrial bioenergetics have different patterns among tissues during aging, we next set out to determine whether impaired ALDH2 function affects mitochondrial metabolism and ALDH2 detox capacity during aging. For that, we used a knock-in mouse carrying an inactivating ALDH2 E504K point mutation (ALDH2^E504K^), which is also present in ~540 million East Asians (17). These animals mimic the human ALDH2 deficiency phenotype (15). We first assessed liver ALDH2 abundance and activity in wild-type, ALDH2^E504K^ heterozygous and homozygous mice (Fig. 2A). Both young and old ALDH2^E504K^ homozygous animals presented a significant decrease in ALDH2 protein levels and catalytic activity compared with wild-type (Figures 2B and C). Notably, aged, but not young, ALDH2^E504K^ heterozygous animals presented a significant decline in liver ALDH2 abundance and activity compared with age-matched wild-type (Fig. 2B-C). Additionally, the aging-dependent increase in liver ALDH2 activity seen in wild-type was lost in ALDH2^E504K^ mice (Fig. 2C). These changes were paralleled by a decrease in ALDH2 levels in old ALDH2^E504K^ animals (Fig. 2B). In summary, aging negatively affects liver ALDH2 levels and function in ALDH2^E504K^ mice; therefore, suggesting a detrimental interaction between ALDH2 deficiency and aging.

**Fig. 2.**
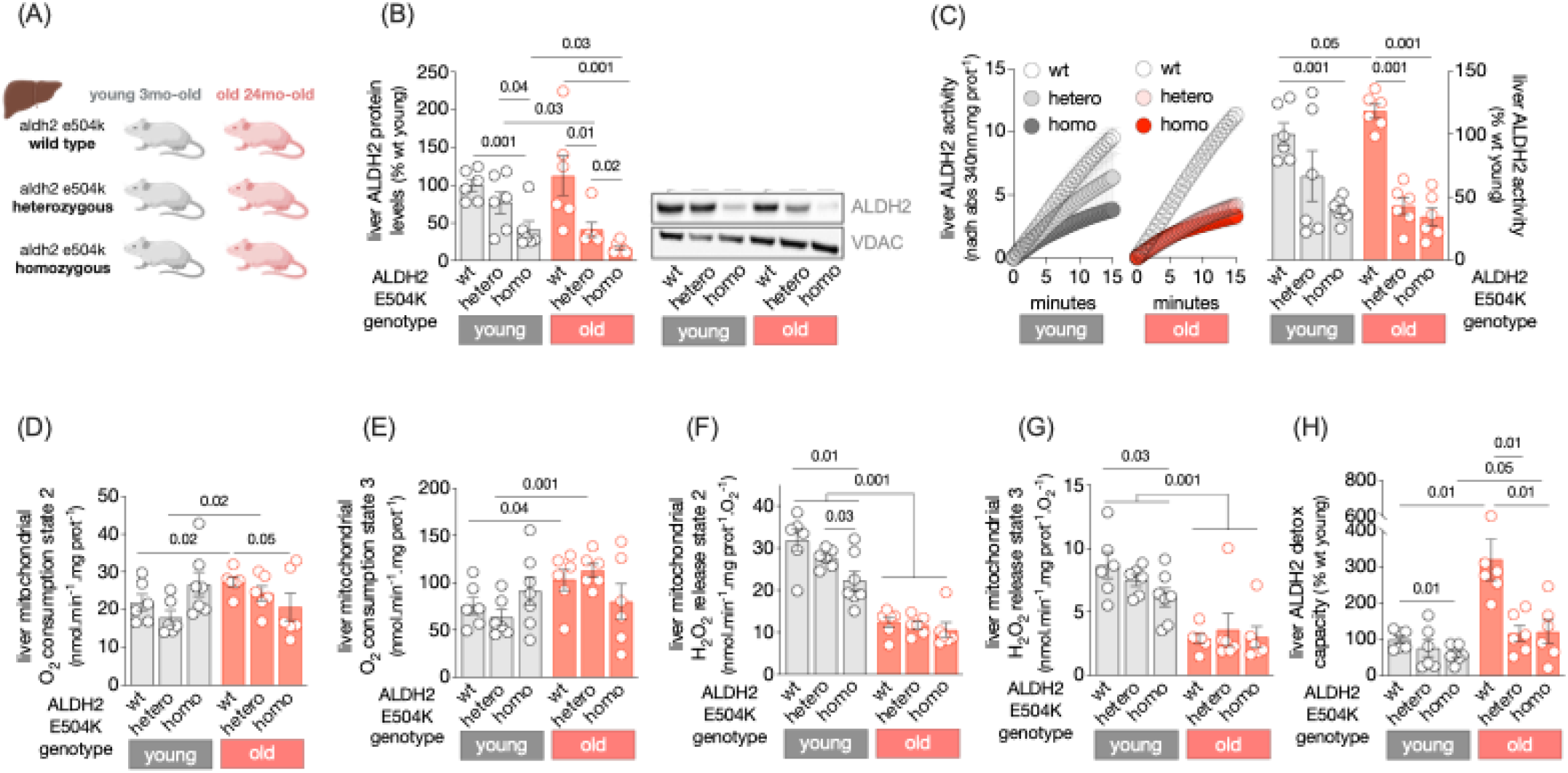
ALDH2^**E504K**^ mutation impairs aging-associated metabolic adaptation in the liver. (A) Schematic panel - mitochondrial metabolism was measured in liver tissue from young (3-month-old, n=6-7) and old (24-month-old, n=6) wild-type, ALDH2^E504K^ heterozygous and ALDH2^E504K^ homozygous male C57BL/6J mice. (B) ALDH2 protein levels (left panel: quantification - individual data and average of all samples, normalized by VDAC; right panel: representative ALDH2 and VDAC immunoblots). (C) ALDH2 activity (left panel: average slope; right panel: quantification - individual data and average of all samples, expressed as % of young wild-type mice), (D–E) mitochondrial O_2_ consumption and (F–G) mitochondrial H_2_O_2_ release (expressed as H_2_O_2_/O_2_) measured in isolated mitochondria under state 2 (after addition of succinate 10 mM, malate 2 mM, glutamate 10 mM) and state 3 (after addition of ADP 4 mM) respiratory rates, and (H) ALDH2 detox capacity (ALDH2 activity divided by mitochondrial H_2_O_2_ release, expressed as % of young wild-type mice) in liver tissue from young (3-month-old) and old (24-month-old) wild-type (wt), ALDH2^E504K^ heterozygous (hetero), and ALDH2^E504K^ homozygous (homo) mice (n=6-7 per group). Data are presented as individual values and means ± SEM. Statistical significance (p<0.05) was assessed using two-way analysis of variance (ANOVA) followed by Tukey’s post-hoc test. P values are individually shown in each panel.

Next, we tested whether impaired liver aldehyde metabolism in ALDH2^E504K^ mice affect mitochondrial function during aging. The aging-related increase in liver mitochondrial oxygen consumption [state 2 and state 3 respiratory rates] seen in wild-type was lost in ALDH2^E504K^ homozygous, but not in heterozygous, mice (Fig. 2D-E). Wild-type and heterozygous ALDH2^E504K^ animals presented similar mitochondrial respiratory rates at both ages (Fig. 2D-E). However, during aging, ALDH2^E504K^ homozygous mice displayed a significant decline in liver mitochondrial oxygen consumption compared with age-matched wild-type (Fig. 2D-E). We next measured the effect of inactivating ALDH2 E504K point mutation on liver mitochondrial oxidative stress. Young ALDH2^E504K^ homozygous, but not heterozygous, mice displayed decreased state 2 and state 3 hydrogen peroxide release compared with age-matched wild-type (Fig. 2F-G). During aging, there was a similar decline in liver mitochondrial state 2 and state 3 hydrogen peroxide release in all genotypes compared with young animals (Fig. 2F-G). Strikingly, the improved liver ALDH2 detox capacity seen in aged wild-type animals was completely lost in ALDH2^E504K^ mice (Fig. 2H).

### 3.3 Cardiac ALDH2 buffering capacity is compromised in both young and aged ALDH2 deficient mice

We next assessed cardiac ALDH2 abundance and activity in wild-type and ALDH2^E504K^ mice during aging (Fig. 3A). ALDH2^E504K^ heterozygous and homozygous mice displayed a genotype-dependent decline in ALDH2 protein levels and catalytic activity at both ages compared with wild-type (Figures 3B and C). Of interest, only ALDH2^E504K^ heterozygous animals presented an aging-dependent decline in cardiac ALDH2 abundance (Fig. 3B-C). All these changes in cardiac ALDH2 abundance and catalytic activity described in young and old ALDH2^E504K^ mice were accompanied by slight modifications in mitochondrial bioenergetics. Aging-related decline in state 2 (basal) mitochondrial respiratory rates seen in wild-type was abolished in ALDH2^E504K^ heterozygous mice (Fig. 3D). Similarly, aging-associated increase in state 3 (ADP-coupled) mitochondrial respiratory rates seen in wild-type was lost in ALDH2^E504K^ homozygous mice (Fig. 3E). Young ALDH2^E504K^ heterozygous and homozygous mice presented elevated state 2 (basal) hydrogen peroxide release compared with age-matched wild type (Fig. 3F). Similar increase in hydrogen peroxide release was observed in ADP-stimulated mitochondria from old ALDH2^E504K^ homozygous mice versus age-matched controls (Fig. 3G). As consequence, disrupted ALDH2 function and mitochondrial redox balance compromised cardiac ALDH2 detox capacity in both young and aged ALDH2 ^E504K^ mice compared with wild-type (Fig. 3H).

**Fig. 3.**
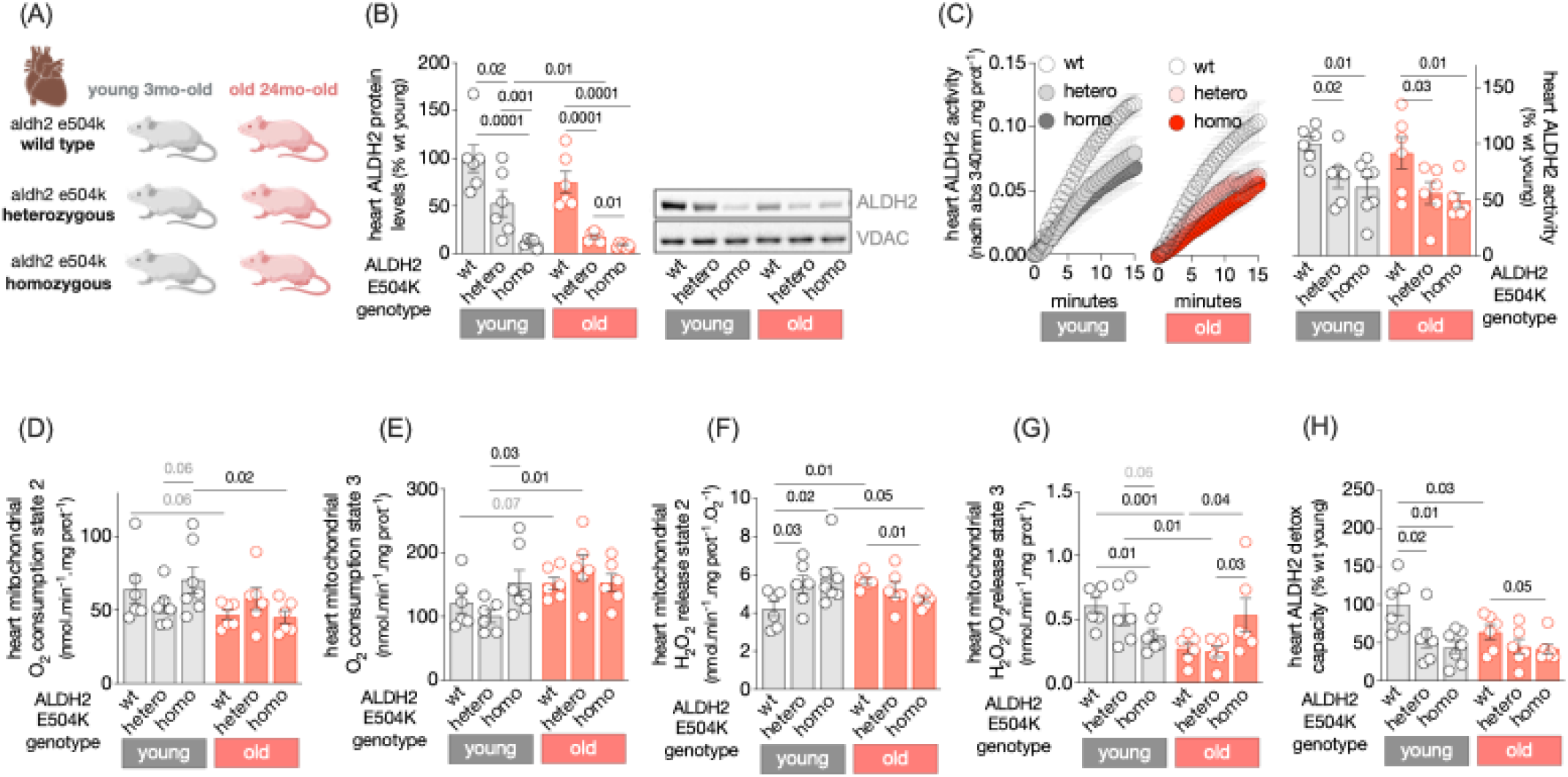
Cardiac ALDH2 activity and mitochondrial bioenergetics in aging ALDH2^**E504K**^ mice. (A) Schematic panel - mitochondrial metabolism was measured in cardiac tissue from young (3-month-old, n=6-7) and old (24-month-old, n=6) wild-type, ALDH2^E504K^ heterozygous and ALDH2^E504K^ homozygous male C57BL/6J mice. (B) ALDH2 protein levels (left panel: quantification - individual data and average of all samples, normalized by VDAC; right panel: representative ALDH2 and VDAC immunoblots). (C) ALDH2 activity (left panel: average slope; right panel: quantification - individual data and average of all samples, expressed as % of young wild-type mice), (D–E) mitochondrial O_2_ consumption and (F–G) mitochondrial H_2_O_2_ release (expressed as H_2_O_2_/O_2_) measured in isolated mitochondria under state 2 (after addition of succinate 10 mM, malate 2 mM, glutamate 10 mM) and state 3 (after addition of ADP 4 mM) respiratory rates, and (H) ALDH2 detox capacity (ALDH2 activity divided by mitochondrial H_2_O_2_ release, expressed as % of young wild-type mice) in cardiac tissue from young (3-month-old) and old (24-month-old) wild-type (wt), ALDH2^E504K^ heterozygous (hetero), and ALDH2^E504K^ homozygous (homo) mice (n=6-7 per group). Data are presented as individual values and means ± SEM. Statistical significance (p<0.05) was assessed using two-way analysis of variance (ANOVA) followed by Tukey’s post-hoc test. P values are individually shown in each panel.

### 3.4 Brain ALDH2 buffering capacity is decreased in both young and aged ALDH2 deficient mice

Here we determined the degree of changes in brain ALDH2 levels and activity in wild-type and ALDH2^E504K^ mice during aging (Fig. 4A). ALDH2^E504K^ mice displayed a genotype-dependent decline in ALDH2 protein levels and catalytic activity at both ages compared with age-matched wild-type (Fig. 4B-C). The age-related accumulation of ALDH2 protein levels seen in wild-type was lost in ALDH2^E504K^ mice (Figures 4B). Of interest, all animals, including wild-type and ALDH2^E504K^, presented an aging-dependent decline in brain ALDH2 activity (Fig. 4C). We next checked whether these genotype-dependent changes in ALDH2 profile affect mitochondrial oxygen consumption, redox balance and ALDH2 detox capacity during aging. No major changes in brain mitochondrial oxygen consumption and hydrogen peroxide release [state 2 and state 3 respiratory rates] were observed among young animals from different genotypes (Fig. 4D-G). The age-related decrease in brain mitochondrial oxygen consumption and increased hydrogen peroxide release seen in wild-type was lost in ALDH2^E504K^ heterozygous mice (Fig. 4D-G). Notably, all aged animals displayed a significant decrease in brain ALDH2 detox capacity compared with genotype-matched young mice (Fig. 4H). Brain ALDH2 detox capacity was further impaired in both young and old ALDH2^E504K^ homozygous mice compared with age-matched wild-type (Fig. 4H).

**Fig. 4.**
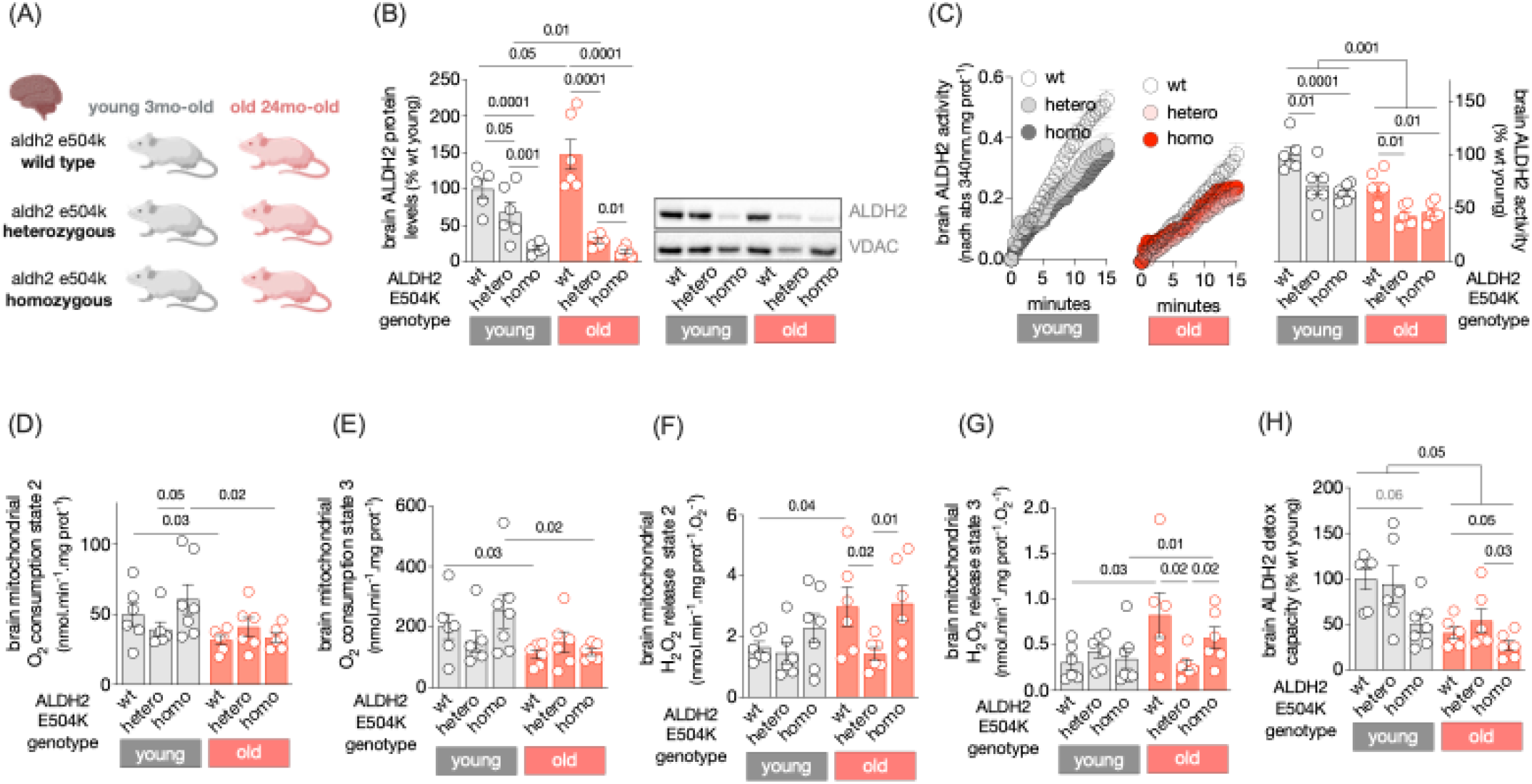
ALDH2^**E504K**^ mutation exacerbates aging-associated mitochondrial dysfunction in the brain. (A) Schematic panel - mitochondrial metabolism was measured in brain tissue from young (3-month-old, n=6-7) and old (24-month-old, n=6) wild-type, ALDH2^E504K^ heterozygous and ALDH2^E504K^ homozygous male C57BL/6J mice. (B) ALDH2 protein levels (left panel: quantification - individual data and average of all samples, normalized by VDAC; right panel: representative ALDH2 and VDAC immunoblots). (C) ALDH2 activity (left panel: average slope; right panel: quantification - individual data and average of all samples, expressed as % of young wild-type mice), (D–E) mitochondrial O_2_ consumption and (F–G) mitochondrial H_2_O_2_ release (expressed as H_2_O_2_/O_2_) measured in isolated mitochondria under state 2 (after addition of succinate 10 mM, malate 2 mM, glutamate 10 mM) and state 3 (after addition of ADP 4 mM) respiratory rates, and (H) ALDH2 detox capacity (ALDH2 activity divided by mitochondrial H_2_O_2_ release, expressed as % of young wild-type mice) in brain tissue from young (3-month-old) and old (24-month-old) wild-type (wt), ALDH2^E504K^ heterozygous (hetero), and ALDH2^E504K^ homozygous (homo) mice (n=6-7 per group). Data are presented as individual values and means ± SEM. Statistical significance (p<0.05) was assessed using two-way analysis of variance (ANOVA) followed by Tukey’s post-hoc test. P values are individually shown in each panel.

### 3.5 Running performance is equally impaired in aged wild-type and ALDH2 deficient mice

Considering all age- and genotype-related changes in mitochondrial bioenergetics and ALDH2 activity in liver, heart and brain from ALDH2^E504K^ animals, we decided to further measure some critical physiological parameters affected by aging (Fig. 5A). First, we assessed body and tissues weights during aging. No major differences in body, heart, lung, liver, kidney and brain weights were observed during aging when comparing age-matched wild-type and ALDH2^E504K^ mice (Fig. 5B and Table S1). Note that all tissues weights were normalized by body weight (Table S1). We next assessed voluntary running performance by placing young and old wild-type and ALDH2^E504K^ animals in cages equipped with running wheels for 14 days. As expected, aged animals displayed a significant decline in both daily and cumulative exercise running distance, time and speed compared with genotype-matched young mice (Fig. 5C-E). No differences in running performance were observed in age-matched wild-type and ALDH2^E504K^ mice (Fig. 5C-E). These findings suggest that tissue-dependent changes in ALDH2 function and mitochondrial bioenergetics seen in ALDH2^E504K^ mice do not affect age-associated modifications in body and tissue weights, and voluntary wheel running performance.

**Fig. 5.**
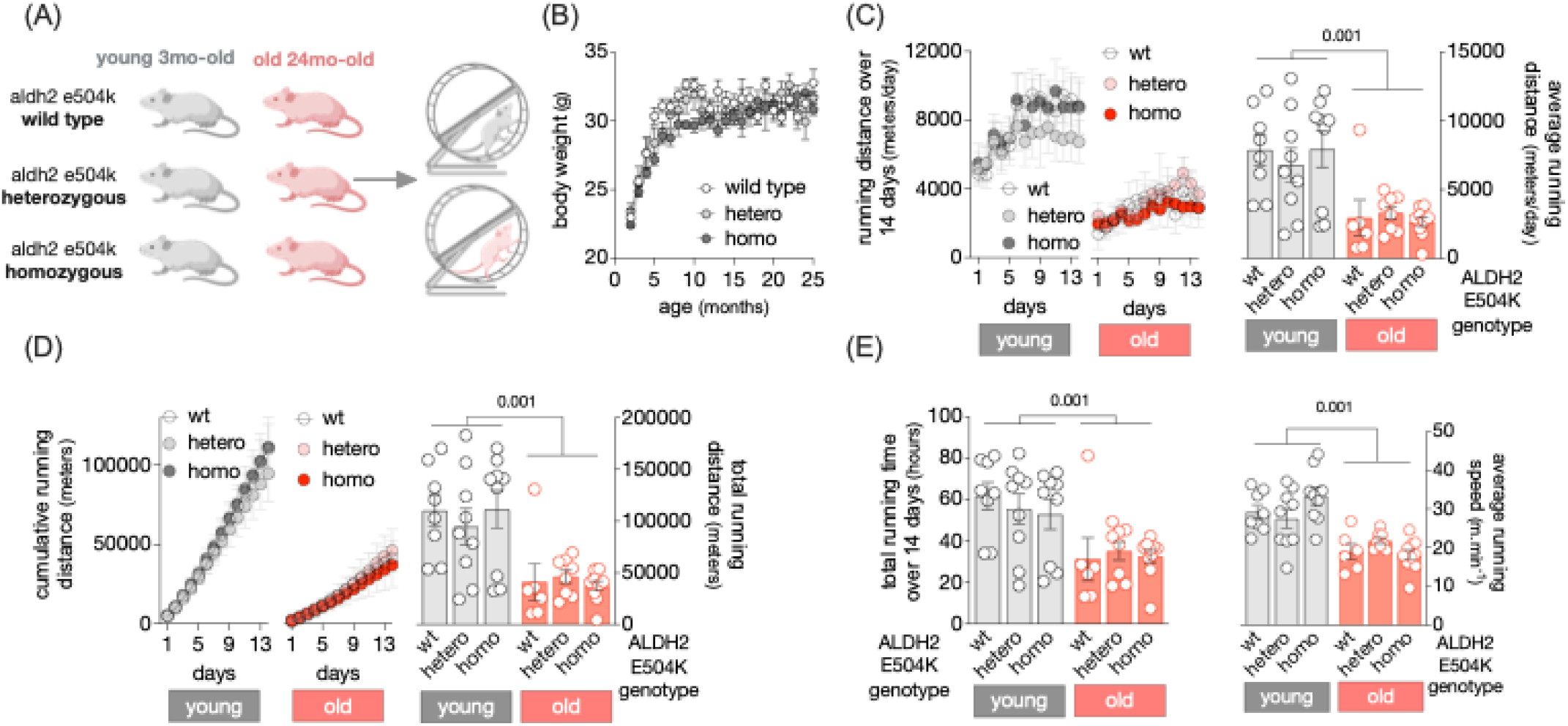
Running performance is equally impaired in aged wild-type and ALDH2 deficient mice. (A) Schematic panel – wheel running performance was measured in young (3-month-old, n=8-9) and old (24-month-old, n=6-9) wild-type, ALDH2^E504K^ heterozygous and ALDH2^E504K^ homozygous male C57BL/6J mice. (B) Body weight over time (g, grams), (C) running distance over 14 days (left panel: daily running distance; right panel: individual data and average running distance over 14 days), (D) cumulative running distance over 14 days (left panel: daily cumulative running distance; right panel: individual data and average cumulative running distance over 14 days), (E) running time and running speed over 14 days (left panel: individual data and average running time over 14 days; right panel: individual data and average running speed over 14 days) in young (3-month-old) and old (24-month-old) wild-type (wt), ALDH2^E504K^ heterozygous (hetero), and ALDH2^E504K^ homozygous (homo) mice (n=6-9 per group). Data are presented as individual values and means ± SEM. Statistical significance (p<0.05) was assessed using two-way analysis of variance (ANOVA) followed by Tukey’s post-hoc test. P values are individually shown in each panel.

### 3.6 Aging-related cognitive and behavioral dysfunction are exacerbated in ALDH2 deficient mice

Here, we set out to perform a series of functional tests to evaluate the impact of aging and/or ALDH2 deficiency in cognitive and behavioral functions in aging mice (Fig. 6A). We decided to run these measurements considering that wild-type aged brain was the only tissue displaying a decline in ALDH2 activity (Fig. 1U), increased mitochondrial oxidative stress (Fig. 1V), and decreased mitochondrial oxygen consumption (Fig. 1W). Note that these metabolic and aging-related changes seen in wild-type brain were exacerbated in ALDH2^E504K^ mice (Fig. 4C and H). To this end, we performed a series of tests to assess anxiety-like behavior (elevated plus maze test), cognitive recognition memory (novel object recognition test), short-term spatial memory (y-maze test) and motor deficits (limb clasping test) (Fig. 6A). In the elevated plus maze test, aged animals presented a significant decline in the amount of time spent in open arms compared with genotype-matched young mice, suggesting an increased anxiety-like behavior during aging (Fig. 6B). The increased anxiety-like behavior in aging mice has been previously described (29). Strikingly, aged ALDH2^E504K^ mice displayed a further decrease in the amount of time spent in open arms compared with age-matched wild-type (Fig. 6B). These findings suggest that ALDH2 deficiency maximizes anxiety-like behavior in aged, but not young, mice.

**Fig. 6.**
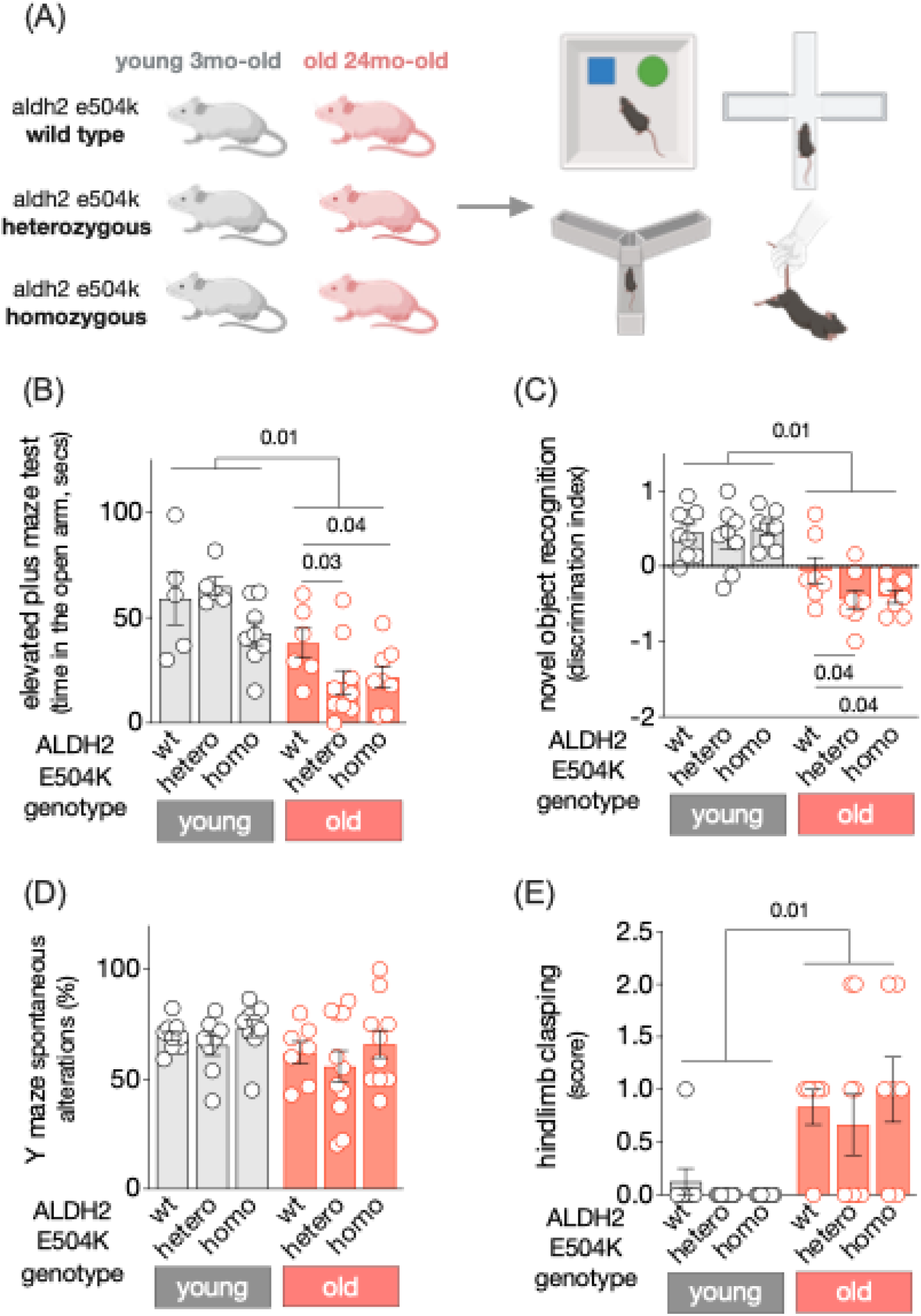
Aging-related cognitive and behavioral dysfunction are exacerbated in ALDH2 deficient mice. (A) Schematic panel – behavioral and cognitive functions were measured in young (3-month-old, n=8-9) and old (24-month-old, n=6-10) wild-type, ALDH2^E504K^ heterozygous and ALDH2^E504K^ homozygous male C57BL/6J mice. (B) Time in seconds spent in the open arm of the elevated plus maze, (C) discrimination index during novel object recognition, (D) Y-maze spontaneous alternations (%), and (E) hindlimb clasping score (0–4 scale) in young (3-month-old) and old (24-month-old) wild-type (wt), ALDH2^E504K^ heterozygous (hetero), and ALDH2^E504K^ homozygous (homo) mice (n=6-10 per group). Data are presented as individual values and means ± SEM. Statistical significance (p<0.05) was assessed using two-way analysis of variance (ANOVA) followed by Tukey’s post-hoc test. P values are individually shown in each panel.

Next, we performed the novel object recognition test to assess cognitive recognition memory during aging. Aged mice displayed decreased novel object recognition discrimination index when compared with genotype-matched young animals (Fig. 6C). These findings suggest impaired cognitive recognition memory during aging, as previously reported (30). Of interest, aged ALDH2^E504K^ mice had a further decline in cognitive recognition memory compared with age-matched wild-type (Fig. 6C). We did not observe any aging-related changes in short-term spatial memory in wild-type animals, measured by Y maze spontaneous alternations (Figure 6D). These results are similar to findings previously reported elsewhere (31). Additionally, ALDH2^E504K^ mice displayed similar short-term spatial memory when compared with age-matched wild-types (Fig. 6D). We also performed the hindlimb clasping test to assess potential motor deficits in aging mice. Aged animals had higher score in the hindlimb clasping test compared with genotype-matched young animals, suggesting motor impairment or neurological dysfunction in aged mice (Fig. 6E). No changes in hindlimb clasping score were seen when comparing age-matched wild-type and ALDH2^E504K^ mice (Fig. 6E).

Considering the major role of the brain in controlling cognitive and behavioral functions in mice, we decided to perform additional measurements in aging animals including anxiety-like behavior and mobility (open field test); coordination (rotarod test) and body temperature (Fig. 7A). In the open field test, aged mice displayed decreased entries in the center when compared with genotype-matched young animals (Fig. 7B). When assessing the genotype effect, both young and old ALDH2^E504K^ mice presented a decline in entries in the center when compared with age-matched controls (Fig. 7B). Similarly, ALDH2^E504K^ mice showed a reduction in the number of crossing when compared with age-matched wild-type (Fig. 7C). Of interest, aged ALDH2^E504K^ animals, but not wild-type, presented decreased number of crossing when compared with genotype-matched young mice (Fig. 7C). When measuring motor coordination, ALDH2^E504K^ homozygous mice displayed decreased latency to fall in the rotarod test when compared with age-matched wild-type (Fig. 7D). Again, aged ALDH2^E504K^ animals, but not wild-type, presented decreased latency to fall when compared with genotype-matched young mice (Fig. 7D). Finally, no major changes in body temperature were observed among different ages and genotypes (Fig. 7E). Overall, these findings suggest that age and/or ALDH2 deficiency have a detrimental effect on anxiety-like behavior and mobility in mice.

**Fig. 7.**
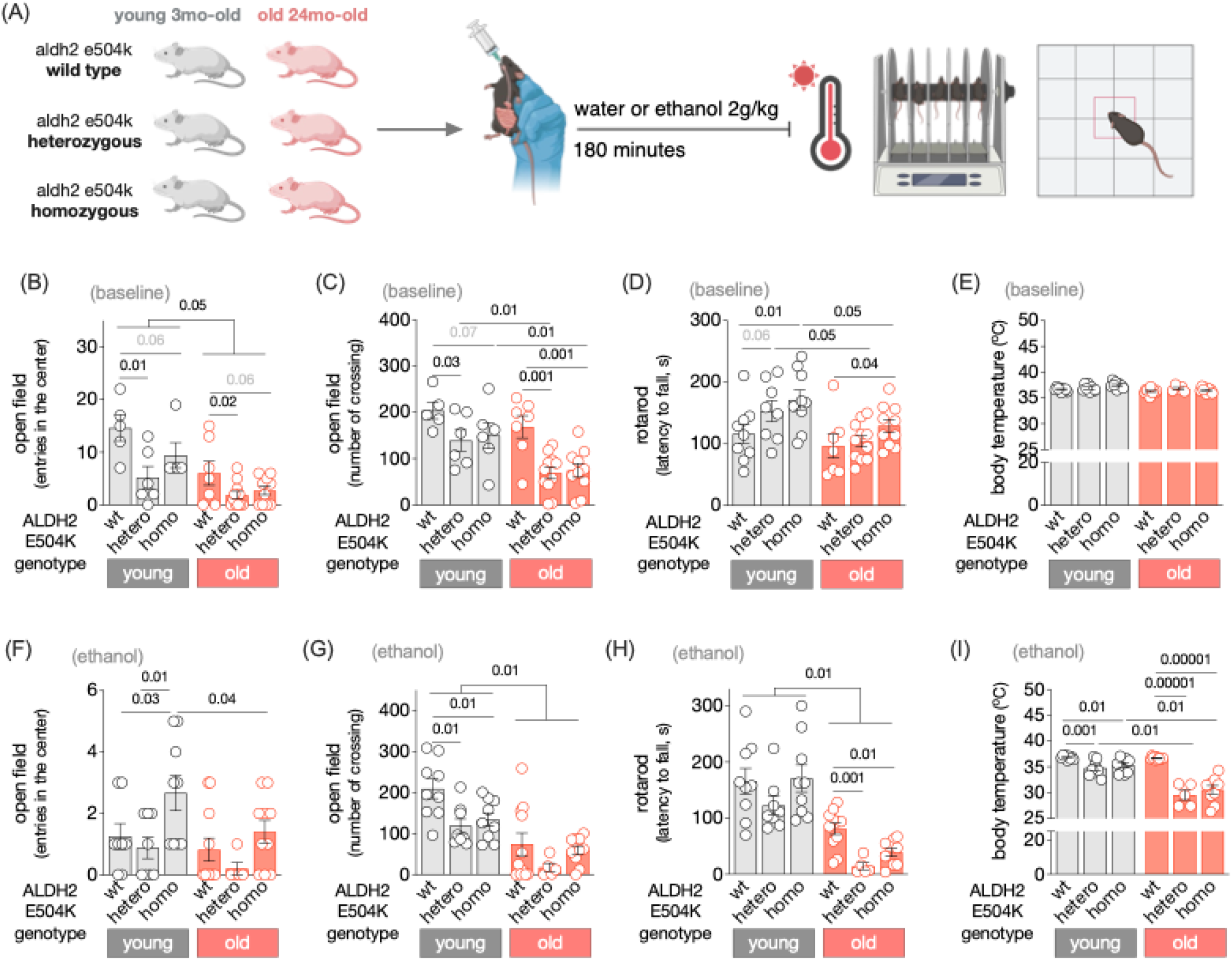
Acute ethanol maximizes aging-related behavioral dysfunction in aging mice. (A) Schematic panel – behavioral functions and body temperature were measured in young (3-month-old, n=5-9) and old (24-month-old, n=6-10) wild-type, ALDH2^E504K^ heterozygous and ALDH2^E504K^ homozygous male C57BL/6J mice 180 minutes after administration of vehicle or ethanol (2g/kg, gavage). (B) Open field test (number of entries in the center), (C) Open field test (number of crossing), (D) rotarod test (latency to fall in seconds) and (E) body temperature 180 minutes after administration of vehicle (baseline). (F) Open field test (number of entries in the center), (G) Open field test (number of crossing), (H) rotarod test (latency to fall in seconds) and (I) body temperature 180 minutes after administration of ethanol (2g/kg, gavage). All measurements were performed in young (3-month-old) and old (24-month-old) wild-type (wt), ALDH2^E504K^ heterozygous (hetero), and ALDH2^E504K^ homozygous (homo) mice (n=5-10 per group). Data are presented as individual values and means ± SEM. Statistical significance (p<0.05) was assessed using two-way analysis of variance (ANOVA) followed by Tukey’s post-hoc test. P values are individually shown in each panel.

We next set out to explore whether increasing aldehydic load acutely, by administering ethanol (2g/kg, equivalent to binge drinking in humans), perturb age- and/or genotype-related behaviors in mice (Fig. 7A). This complementary experimental strategy is needed considering the impaired aldehyde metabolism in aging brain (Fig. 1), as well as the negative impact of ALDH2 on cognitive and behavioral functions in aging mice (Fig. 6). No major differences of entries in the center (open field test) were seen between wild-type and ALDH2^E504K^ heterozygous mice after acute ethanol challenge (Fig. 7F). Ethanol administration decreased the number of crossing (open filed test) and the latency to fall (rotarod test) in aged animals from all genotypes when compared with genotype-matched young mice (Fig. 7G-H). Strikingly, old ALDH2^E504K^ mice displayed a further decline in the latency to fall (rotarod test) when compared with age-matched wild-type after ethanol exposure (Fig. 7H). Finally, ethanol administration caused a decline in body temperature in both young and old ALDH2^E504K^ animals compared with age-matched wild-type (Fig. 7I). Of interest, aged ALDH2^E504K^ animals, but not wild-type, exposed to ethanol presented decreased body temperature when compared with genotype-matched young mice (Fig. 7I). In summary, acute ethanol challenge potentiated the aging effect on reduced motor coordination and body temperature in ALDH2 deficient mice.

### 3.7 Impact of aging and ALDH2 deficiency on ALDH2 response to selective pharmacological modulators

We and others have reported the benefits of pharmacological ALDH2 activation in animal models of age-related disorders including myocardial infarction (14), heart failure (5) and Alzheimer disease (32). However, these studies were all conducted using young adult animals; therefore, excluding the aging confounding effect. Here, we decided to evaluate the individual and combined effect of aging and ALDH2 deficiency on ALDH2 response to selective pharmacological activation [AD-5591 (13)] and inhibition [cvt-10216, (7)]. For that, we isolated mitochondria from liver, heart and brain of wild-type and ALDH2^E504K^ aging animals, individually incubated them with ALDH2 modulators *in vitro* and measured ALDH2 catalytic activity. *In vitro* incubation of liver mitochondria with AD-5591 increased ALDH2 catalytic activity compared with vehicle in all ages and genotypes studied (Fig. 8A and S2A). However, the degree of AD-5591-mediated ALDH2 activation in both young and old ALDH2^E504K^ homozygous was significantly smaller compared with the other genotypes (Fig. 8A). Similarly, the cvt-10216 compound decreased ALDH2 activity in wild-type and ALDH2^E504K^ heterozygous at both ages compared with vehicle (Fig. 8A and S2A). ALDH2^E504K^ homozygous did not respond to cvt-10216 inhibition (Fig. 8A and S2A).

**Fig. 8.**
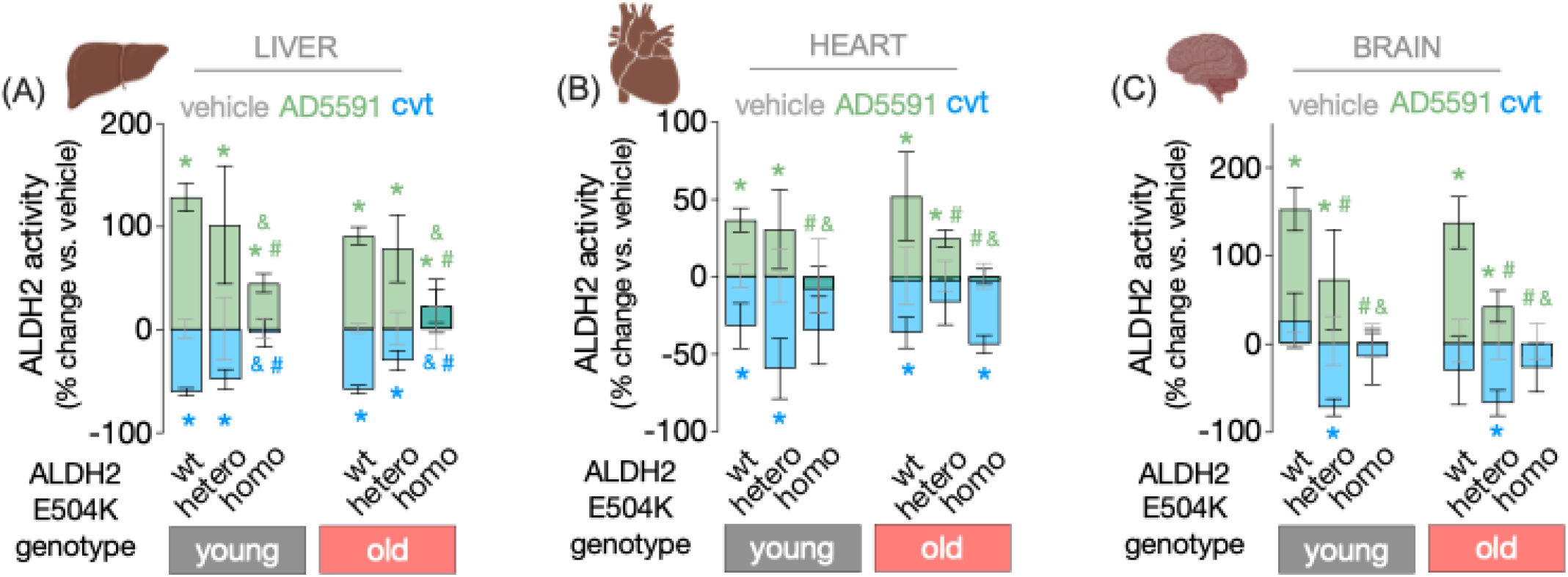
Impact of aging and ALDH2 deficiency on ALDH2 response to selective pharmacological modulators. Tissue-specific ALDH2 activity after *in vitro* incubation with vehicle (DMSO), AD-5591 (1 µM), or cvt-10216 (1 µM) for 10 min at 37 °C. (A) Liver, (B) brain, and (C) heart tissue lysates from young (3-month-old) and old (24-month-old) wild-type (wt), heterozygous ALDH2^E504K^ (hetero) and homozygous ALDH2^E504K^ (homo) male C57BL/6J mice (n = 6 per group) were incubated with ALDH2 modulators (AD-5591 and cvt-10216. Data are presented as means ± SEM. Statistical significance (p<0.05) was assessed using two-way analysis of variance (ANOVA) followed by Tukey’s post-hoc test. * p<0.05 vs. vehicle, # p<0.05 vs. age-matched wild-type, and & p<0.05 vs. age-matched heterozygous ALDH2^E504K^.

Next, we assessed ALDH2 response to AD-5591 and cvt-10216 compounds in isolated cardiac mitochondria. *In vitro* incubation of heart mitochondria with AD-5591 enhanced ALDH2 activity in wild-type and ALDH2^E504K^ heterozygous at both ages compared with vehicle (Fig. 8B and S2B). Note that old, but not young, ALDH2^E504K^ heterozygous mice displayed a lower magnitude of AD-5591-induced increase in ALDH2 activity compared with age-matched wild-type. ALDH2^E504K^ homozygous did not respond to AD-5591 at both ages (Fig. 8B and S2B). The cvt-10216 compound decreased cardiac ALDH2 catalytic activity in wild-type at both ages (Fig. 8B and S2B). Cardiac mitochondria from young ALDH2^E504K^ heterozygous and old ALDH2^E504K^ homozygous also responded to cvt-10216 (Fig. 8B and S2B).

Finally, we determined whether brain mitochondria respond to pharmacological ALDH2 modulators. *In vitro* incubation of brain mitochondria with AD-5591 boosted ALDH2 activity in wild-type and ALDH2^E504K^ heterozygous at both ages compared with vehicle (Figures 8C and S2C). However, the amplitude of ALDH2 activation in ALDH2^E504K^ heterozygous brain was significantly smaller at both ages compared with wild-type (Figure 8C and S2C). Brain mitochondria from ALDH2^E504K^ homozygous mice did not respond to AD-5591 at both ages (Figure 8C and S2C). Intriguingly, only mitochondria isolated from ALDH2^E504K^ heterozygous brain was responsive to cvt-10216-induced ALDH2 inhibition (Figure 8C and S2C). Overall, these findings demonstrate that ALDH2 catalytic activity in mitochondria isolated from liver, heart and brain respond similarly to AD-5591 when comparing young and old wild-type animals. A similar response is observed in liver and heart tissues exposed to cvt-10216. Of interest, both age and ALDH2 genotype synergistically affected ALDH2 response to AD-5591 and cvt-10216 *in vitro*.

## 4. DISCUSSION

Age-related diseases result from chronic exposure to genetic and/or environmental factors, culminating in the progressive and irreversible degeneration of cellular functions, tissues, and ultimately the organism as a whole (33). As such, identifying clinically relevant molecular events in aging remains a major priority toward the development of more effective interventions to treat age-related diseases and improve healthy aging. As an effort for the identification and validation of relevant metabolic pathways in aging, our findings provide a comprehensive analysis of how ALDH2 metabolism and mitochondrial bioenergetics are modulated and integrated across different tissues during aging, and how these processes are further influenced by the presence of the highly prevalent human inactivating ALDH2 E504K point mutation, present in approximately 8% of the world population (~560 million people) (17).

During aging, there is an increase in circulating levels of reactive aldehydes, which readily form covalent adducts with proteins, lipids and nucleic acids (34). Some of these aldehydes such as acetaldehyde, 4-hydroxynonenal and malondialdehyde are produced within mitochondria during lipid peroxidation (13), ethanol metabolism (7) and excessive glucose metabolism (6). Under physiological conditions, these mitochondrially-derived aldehydes are rapidly metabolized into less reactive molecules by ALDH2; therefore, neutralizing their toxicity. However, accumulation of these toxic aldehydes due to either its overproduction or impaired turnover is sufficient to cause mitochondrial dysfunction (5), disrupt cellular homeostasis (13) and contribute to the establishment and progression of several age-related diseases (25, 32, 35). Under this scenario, the efficiency of mitochondrial aldehyde metabolism becomes a limiting factor for mitochondrial and tissue health during aging (2).

Considering that aldehyde metabolism is directly linked to mitochondrial respiration, it is expected that tissue-specific heterogeneity in aldehyde metabolism will locally impact mitochondrial aldehydic load and bioenergetics during aging. Here, we show that liver mitochondrial metabolism is relatively resilient to aging, at least in part, due to its greater aldehydic detox capacity compared with heart and brain tissues. As proof of concept, genetic ALDH2 deficiency is sufficient to abolish the aging-related improvements in liver aldehydic detox capacity and mitochondrial bioenergetics. Aging-induced liver mitochondrial hypermetabolism and resilience have been attributed, at least in part, to increased mitochondrial proton leakage (36, 37). Consistent with this understanding, our results indicate that aged liver exhibits mild mitochondrial uncoupling, as evidenced by elevated mitochondrial oxygen consumption accompanied by decreased mitochondrial hydrogen peroxide release.

Aging is among the strongest predictors of cardiovascular and neurological health in humans (38, 39). In cardiac and brain tissues, we observed a decline in aldehydic detox capacity during aging compared with liver; therefore, favoring aldehydic load and mitochondrial dysfunction. It is important to note that cardiac and brain tissues exhibit the highest energetic demand in the body (40). These observations suggest that decreased aldehydic detox capacity along with elevated mitochondrial oxygen consumption make cardiac and brain tissues more susceptible (versus liver) to mitochondrial dysfunction and aldehydic load during aging. We have previously reported that increasing aldehyde clearance through pharmacological activation of ALDH2 is sufficient to improve mitochondrial dysfunction (5) and treat age-related cardiovascular diseases in rodents (13, 35); therefore, supporting an ALDH2 mechanism for mitochondrial dysfunction in degenerative diseases.

Here we show that ALDH2 detox capacity and mitochondrial bioenergetics are largely compromised in brain tissue during aging. These findings are reinforced by our loss-of-function approach, where genetic ALDH2 deficiency maximizes aging-related cognitive and behavioral dysfunction in mice. Aldehydic load has been associated with neurological disorders in both humans (41) and mice (42). A recent meta-analysis correlated ALDH2^E504K^ mutation and Alzheimer disease development (20). ALDH2 knockout mice display exacerbated cognitive impairment and accumulation of Alzheimer disease biomarkers (43). Additionally, ethanol administration-induced neuroinflammation is maximized in young ALDH2^E504K^ mice (32). Here we report that aging potentiates ethanol-induced behavior dysfunction, having a major effect in ALDH2^E504K^ mice. Strikingly, selective ALDH2 activation was sufficient to mitigates ethanol-induced neuroinflammation and Alzheimer disease-associated pathology in young mice (32).

Together, our findings reveal how ALDH2 metabolism and mitochondrial bioenergetics are coordinated across tissues during aging, and how these processes are affected by the common inactivating ALDH2 E504K mutation. The liver metabolism was relatively resilient to aging, while heart and brain tissues displayed impaired ALDH2 activity and dysfunctional mitochondrial bioenergetics during aging. Notably, these changes in aging brain metabolism were followed by impaired cognitive and behavioral functions in wild-type mice, which were maximized by the presence of the ALDH2 E504K mutation or after ethanol challenge. These are known experimental conditions that induce aldehydic load (7). Finally, we provide evidence that AD-5591 boost *in vitro* ALDH2 activity in liver, heart and brain tissues from aging wild-type and ALDH2^E504K^ heterozygous mice. These findings provide a framework to guide future investigations using AD-5591 in aged animals, which display different aldehyde detox activity and mitochondrial bioenergetics compared with young organism. We have previously reported the efficacy of AD-5591 to treat heart failure in young adult rats (13). Further studies in aging animal models and humans are needed to establish AD-5591 as an effective therapy for aging-related diseases. It is important to highlight that a multiple-dose, phase 1 clinical trial evaluating the safety of a small molecule activator of ALDH2 was recently completed with no major side effects reported in humans (NCT03686930); therefore, supporting the use of this class of ALDH2 activators in humans.

## FIGURE LEGENDS

**Fig. S1.**
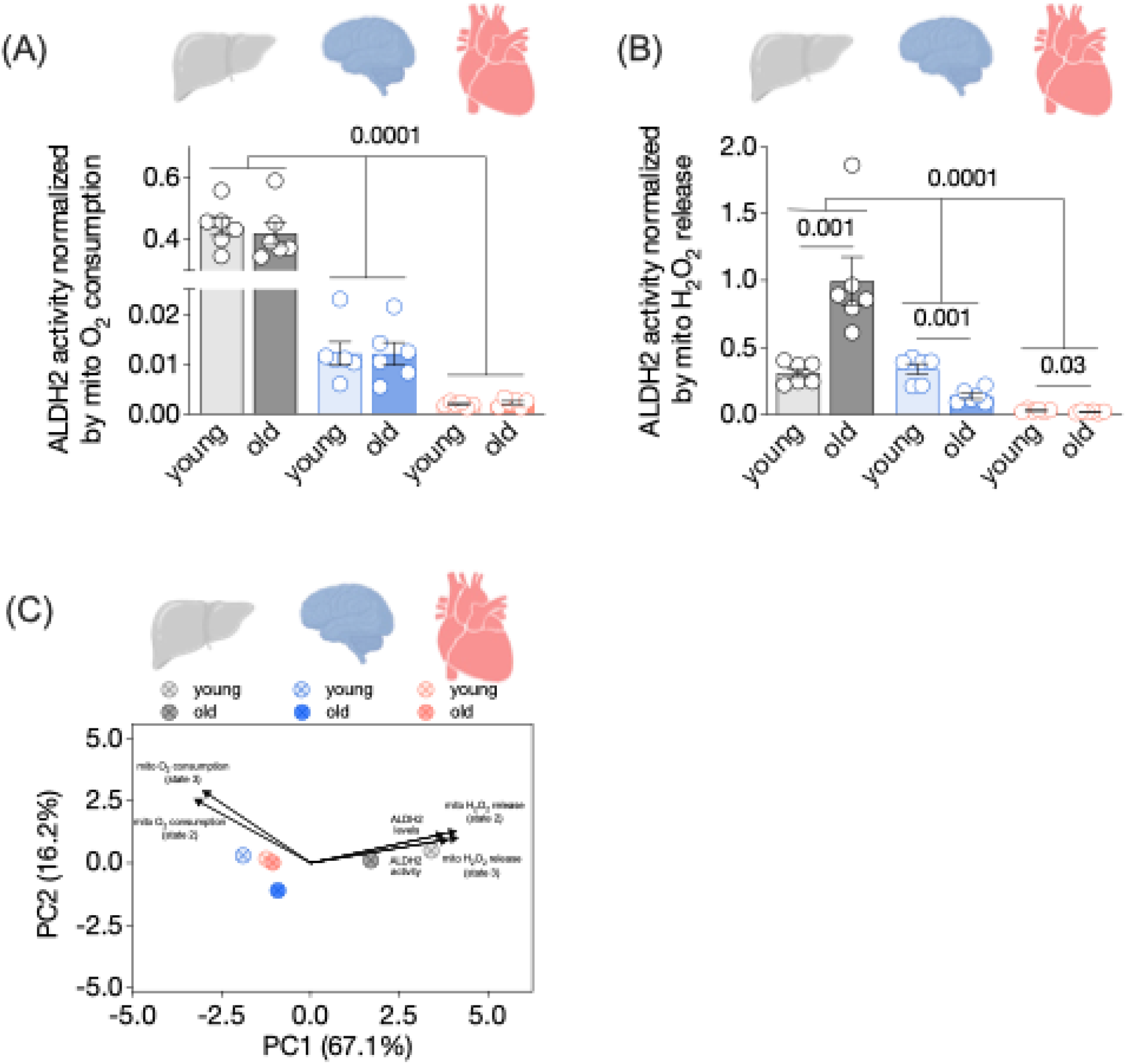
Tissue-specific ALDH2 activity and mitochondrial bioenergetics during aging. (A) ALDH2 activity normalized by state 2 mitochondrial O_2_ consumption and (B) ALDH2 activity normalized by state 2 H_2_O_2_ release in liver, brain, and heart from young (3-month-old) and old (24-month-old) wild-type male C57BL/6J mice (n=6 per group). (C) Principal component analysis (PCA) loading vectors and centroids used to provide biological interpretation of the principal components (related to Fig. 1X) in liver, brain, and heart from young (3-month-old) and old (24-month-old) wild-type male C57BL/6J mice (n=6 per group). PC1 was primarily associated with variables related to ALDH2 activity and hydrogen peroxide release, whereas PC2 was more closely associated with mitochondrial respiratory parameters. Data are presented as individual values and means ± SEM. Statistical significance (p<0.05) was assessed using two-way analysis of variance (ANOVA) followed by Tukey’s post-hoc test. P values are individually shown in each panel.

**Fig. S2.**
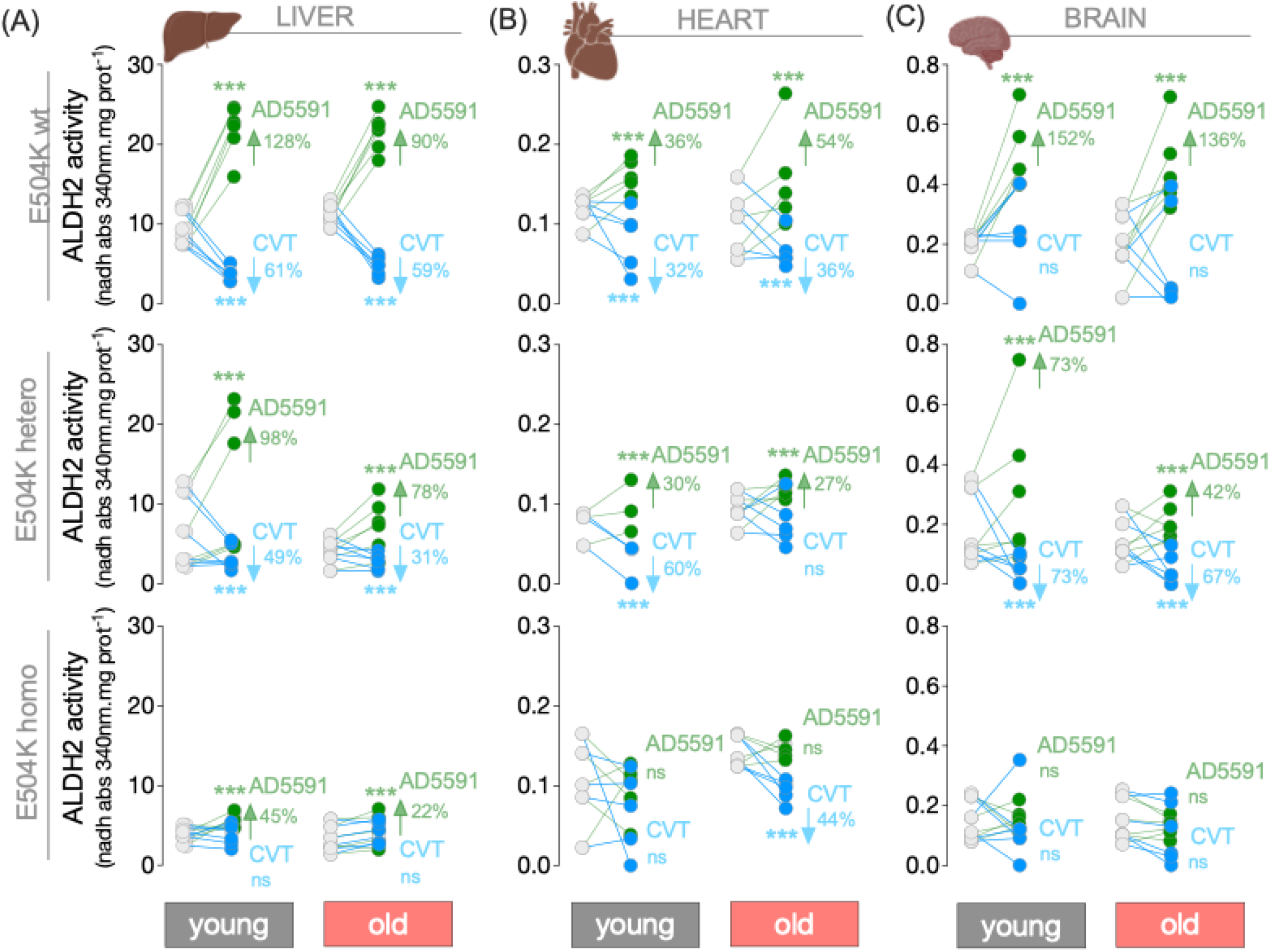
Tissue-specific pharmacological modulation of ALDH2 activity across aging and ALDH2^**E504K**^ genotypes. Tissue-specific ALDH2 activity after *in vitro* incubation with vehicle (DMSO), AD-5591 (1 µM), or cvt-10216 (1 µM) for 10 min at 37 °C. (A) Liver, (B) brain, and (C) heart tissue lysates from young (3-month-old) and old (24-month-old) wild-type (wt), heterozygous ALDH2^E504K^ (hetero), and homozygous ALDH2^E504K^ (homo) male C57BL/6J mice (n = 6 per group) were incubated with ALDH2 modulators (AD-5591 and cvt-10216. Data are presented as individual values and means ± SEM. Statistical significance (P < 0.05) was assessed using two-tailed Student’s t-tests for pairwise comparisons. *** p<0.05 vs. age-matched vehicle.

**Supplementary Table 1.** Physiologic parameters of young and old male wdd type, heterozygous and homozygous knock-in mice carrying the ALDH2 E504K mutation.

| Parameter | young (3mo-old) |  |  | old (24mo-old) |  |  |
| --- | --- | --- | --- | --- | --- | --- |
|  | wt | ALDH2 hetero | ALDH2 homo | wt | ALDH2 hetero | ALDH2 homo |
| BW (g) | 26.45±0.64 | 26.19±0.31 | 24.12±0.58 | 31.73±1.32<sup>‡</sup> | 33.06±0.53<sup>‡</sup> | 33.01±0.71<sup>‡</sup> |
| HW (g) | 0.14±0.01 | 0.14±0.01 | 0.14±0.01 | 0.21±0.01<sup>‡</sup> | 0.21±0.01<sup>‡</sup> | 0.19±0.01<sup>‡</sup> |
| HW/BW (mg/g) | 5.33±0.1 | 5.51±0.17 | 5.95±0.58 | 6.64±0.40 | 6.03±0.25 | 5.79±0.26 |
| LuW (g) | 0.16±0.00 | 0.16±0.01 | 0.16±0.01 | 0.20±0.01 | 0.23±0.02<sup>‡</sup> | 0.22±0.01 |
| LuW/BW (mg/g) | 1116.16±27.9 | 1139.01±73.84 | 1135.51±67.95 | 975.85±60.84 | 1117.16±77.06 | 1183.70±82.24 |
| LiW (g) | 1.43±0.03 | 1.35±0.05 | 1.41±0.07 | 1.72±0.10 | 1.66±0.11 | 1.55±0.06 |
| LiW/BW (mg/g) | 54.07±1.48 | 51.43±1.69 | 58.29±1.90 | 54.17±1.73 | 47.99±3.50 | 46.36±1.27 |
| KiW (g) | 0.17±0.00 | 0.17±0.01 | 0.17±0.01 | 0.22±0.01 | 0.25±0.02<sup>‡</sup> | 0.23±0.01 |
| KiW/BW (mg/g) | 119.40±0.90 | 130.02±4.62 | 118.60±5.77 | 127.84±6.09 | 161.22±18.10 | 148.14±5.06 |
| BrW (g) | 0.31±0.01 | 0.31±0.01 | 0.31±0.01 | 0.42±0.01<sup>‡</sup> | 0.42±0.01<sup>‡</sup> | 0.41±0.01<sup>‡</sup> |
| BrW/BW (mg/g) | 2171.58±120.46 | 2191.71±88.65 | 2214.31±97.56 | 2021.48±102.19 | 2027.98±74.17 | 2179.67±147.74 |
| TAW (g) | 0.04±0.00 | 0.04±0.00 | 0.04±0.00 | 0.04±0.00 | 0.04±0.00 | 0.04±0.00 |
| TAW/BW (mg/g) | 1.56±0.04 | 1.60±0.01 | 1.53±0.06 | 1.36±0.04 | 1.24±0.04<sup>‡</sup> | 1.31±0.06<sup>‡</sup> |

Body weight (BW), heart weight (HW), heart weight/body weight (HW/BW), lung weight (LuW), lung weight/body weight (LuW/BW), liver weight (LiW), liver weight/body weight (LiW/BW), kidney weight (KiW), kidney weight/body weight (KiW/BW), brain weight (BrW), brain weight/body weight (BrW/BW), tibialis anterior weight (TAW), tibialis anterior weight/body weight (TAW/BW) in young and old wild type, heterozygous and homozygous knock-in mice carrying the
ALDH2 E504K mutation (n=6-14). Data were analyzed by two-way ANOVA. Error bars indicate SEM. <sup>‡</sup>P<0.05 vs young.

